# High-level ceftazidime-avibactam resistance in *Escherichia coli* conferred by the novel plasmid-mediated beta-lactamase CMY-185 variant

**DOI:** 10.1101/2023.02.03.527067

**Authors:** William C. Shropshire, Bradley T. Endres, Jovan Borjan, Samuel L. Aitken, William C. Bachman, Christi L. McElheny, Ayesha Khan, Micah M. Bhatti, Pranoti Saharasbhojane, Akito Kawai, Ryan K. Shields, Samuel A. Shelburne, Yohei Doi

## Abstract

**Objectives:** To characterize a *bla*_CMY_ variant associated with ceftazidime-avibactam (CZA) resistance from a serially collected *Escherichia coli* isolate.

**Methods:** A patient with an intra-abdominal infection due to recurrent *E. coli* was treated with CZA. On day 48 of CZA therapy, *E. coli* with a CZA MIC of >256 mg/L was identified from abdominal drainage. Illumina WGS was performed on all isolates to identify potential resistance mechanisms. Site-directed mutants of CMY β-lactamase were constructed to identify amino acid residues responsible for CZA resistance.

**Results:** WGS revealed that all three isolates were *E. coli* ST410. The CZA-resistant strain uniquely acquired a novel CMY β-lactamase gene, herein called *bla*_CMY-185_, harbored on an IncIγ-type conjugative plasmid. The CMY-185 enzyme possessed four amino acid substitutions relative to CMY-2 including A114E, Q120K, V211S, and N346Y and conferred high-level CZA resistance with an MIC of 32 mg/L. Single CMY-2 mutants did not confer reduced CZA susceptibility. However, double and triple mutants containing N346Y previously associated with CZA resistance in other AmpC enzymes, conferred CZA MICs ranging between 4 and 32 mg/L as well as reduced susceptibility to the newly developed cephalosporin, cefiderocol. Molecular modelling suggested that the N346Y substitution confers the reduction of avibactam inhibition due to the steric hindrance between the side chain of Y346 and the sulfate group of avibactam.

**Conclusion:** We identified CZA resistance in *E. coli* associated with a novel CMY variant. Unlike other AmpC enzymes, CMY-185 appears to require an additional substitution on top of N346Y to confer CZA resistance.

## Introduction

Avibactam (AVI) is a class A, C and D β-lactamase inhibitor which can improve β-lactam activity against several Gram-negative organisms.^1^ The potent inhibitory profile of AVI has made ceftazidime-avibactam (CZA) one of the preferred therapies for infections caused by *Klebsiella pneumoniae* carbapenemase (KPC)-producing *Enterobacterales*.^2^ CZA is also highly active against non-carbapenemase-producing carbapenem-resistant *Enterobacterales* clinical isolates including *Klebsiella aerogenes*, *Enterobacter cloacae*, and others that produce AmpC or extended-spectrum β-lactamases (ESBLs).^3^

Nevertheless, there are certain AmpC variants that have reduced AVI inhibition. Through a series of experimental passaging studies, a combination of ceftazidime (CAZ) or aztreonam (ATM) with AVI has been shown to select for a variety of amino acid hot spot mutations in derepressed chromosomal AmpC (cAmpC), including N346Y variants found in *Citrobacter freundii* and *Enterobacter cloacae*.^4, 5^ The functional role of the N346Y substitution has been characterized in recombinant, isogenic *E. coli* strains producing cAmpC of *E. cloacae*, PDC-5 (cAmpC of *Pseudomonas aeruginosa)*, and DHA-1 (cAmpC of *Morganella morganii*) with each respective variant conferring increased CZA MICs as a result of reduced AVI activity.^6^ A structural analysis of the AmpC binding pocket with AVI demonstrated that cAmpC N346 was one of eight conserved residues that specifically interact with the sulfate group of AVI.^7^ Furthermore, a cAmpC N346T substitution would be predicted to create steric hindrance with the sulfate group and thus affect AVI binding affinity across multiple cAmpC homologs.^7^

CMY-2 is the most common plasmid-associated AmpC (pAmpC) β-lactamase produced by *Escherichia coli* and other *Enterobacterales* species.^8^ The *bla*_CMY-2_ gene is often detected in association with the mobile IS*Ecp1* element that is likely responsible for its wide transmission across multiple *Enterobacterales* species.^9, 10^ A CMY variant, CMY-172, with K290_V291_A292del, N346I compared with CMY-2, has been reported to confer high-level CZA resistance in *Klebsiella pneumoniae* clinical isolates co-producing KPC-2 and CTX-M-65 in China.^11^

Here, we report serial *E. coli* clinical isolates which developed CZA resistance through the acquisition of a novel CMY-2 variant, CMY-185, which developed a complex set of amino acid mutations during treatment. We confirmed the CZA-resistance correlation with this novel CMY-185 mutant through site-directed mutagenesis of CMY-2 to determine the contribution of single vs combination mutations observed within CMY-185.

## Materials and Methods

### Strains and susceptibility testing

The *E. coli* isolates were identified from a surgical drain of a patient admitted to a hospital in Texas. All three *E. coli* isolates (Ec1 through Ec3) were available for further laboratory evaluation. Initial susceptibility testing was performed with Vitek2 or Etest (bioMérieux, Marcy L’Étoile, France) per routine clinical practice at the hospital laboratory. Resistance was determined using CLSI M100 standards (29^th^ edition).^12^ MICs of the recombinant strains were conducted by the broth microdilution method.

### Whole genome sequencing and computational analyses

Genomic DNA (gDNA) of *E. coli* isolates Ec1, Ec2, and Ec3 was extracted using the QIAGEN DNeasy Blood and tissue kit following manufacturer’s instruction. Library preparation for WGS was completed using the Illumina Nextera kit and sequenced with the Illumina NovaSeq 6000 platform using 150-bp paired-end reads. FastQC-v0.11.9 was used to check read quality upon which SPAdes-v3.15.2 was used to create draft assemblies using the isolate parameter.^13^ Antimicrobial resistance genes were detected using ABRicate-v1.0.0 (Seemann, T. ABRicate GitHub: https://github.com/tseemann/abricate) with the CARD database (Accessed: 2022-02-02).^14^ Plasmid replicon typing was performed with ABRicate-v1.0.0 using the PlasmidFinder database (Accessed: 2022-02-02).^15^ The snippy-multi script (Seemann, T. Snippy GitHub: https://github.com/tseemann/snippy) was used to create a core SNP phylogeny of the study isolates using a previously sequenced ST410 isolate (MB9108) as reference (RefSeq #: GCF_024917735.1). Gubbins-v.3.2.1 was used to create a recombination-free, core genome inferred maximum-likelihood phylogeny.^16^ All ST410 isolates were aligned to a IncIγ *bla*_CMY-42_ positive plasmid, pCMY42-035148 (RefSeq #: GCF_003268695.1), using bwa mem and Samtools-v1.14 to subsequently check breadth of IncIγ coverage to further substantiate PlasmidFinder results. An alignment of FtsI encoding genes from SPAdes assemblies was performed using mafft-v7.505 with mutations called based on comparison to MG1655 reference (Accession #: NC_000913.3). Pairwise SNP distances removing recombination sites were determined using snp-sites-v2.5.1^17^ and snp-dists-v0.8.2 respectively using the gubbins.filtered_polymorphic_sites.fasta file as an input file.

### Construction of CMY mutants

CMY-2 gene was cloned into vector pBCSK(-) using the primers CMY-For XbaI (5’-GCTCTAGACATATGATGAAAAAATCGTTATGCTG-3’) and CMY-Rev BamHI (5’-CGGGATCCTTATTGCAGCTTTTCAAGAATG-3’), which was then electroporated into *E. coli* TOP10 as previously described.^18^ Single, double, and triple mutants of CMY-2 were generated from the CMY-2-encoding vector using Q5 CMY_A114E_For (5’-ACCTATACGGAAGGCGGCCTA-3’) and Q5 CMY_A114E_Rev (5’-GGCTAAGTGCAGCAGGCG-3’), Q5 CMY_Q120K_For (5’-CCTACCGCTGAAGATCCCCGA-3’) and Q5 CMY_Q120K_Rev (5’-CCGCCTGCCGTATAGGTG-3’), Q5 CMY_V211S_For (5’-GCCCGTACACAGTTCTCCGGGACAAC-3’) and Q5 CMY_V211S_Rev (5’-TTCCCTTCGCGATAGCCC-3’), and Q5 CMY_N346Y_For (5’-AAGCTATCCTTACCCTGTCCG-3’) and Q5 CMY_N346Y_Rev (5’-TTGTTTGCCAGCATCACG -3’) and electroporated into *E. coli* TOP10. All mutations were confirmed by Sanger sequencing. **Figure 1** is a flowchart documenting the process by which CMY mutants were generated using CMY-2 as the baseline allele.

**Figure 1.**
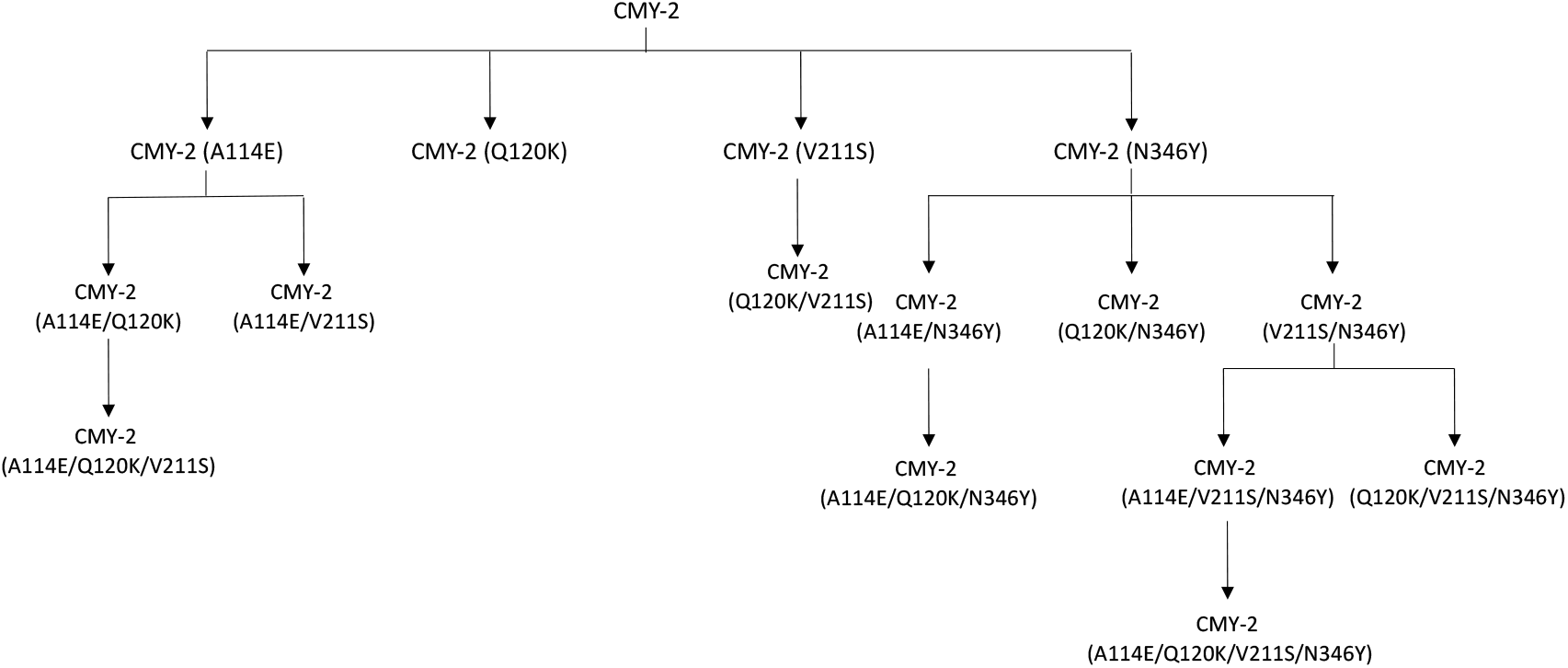
Overview of Site-Directed Mutagenic CMY-2 variants. Parenthetical indicates the amino acid mutation relative to CMY-2. A total of 15 CMY-2 mutants were created using a PCR-based mutagenesis method and subsequently assessed using antimicrobial susceptibility testing.

### Molecular modelling

A homology model of CMY-185 was generated using the CMY-136 structure (PBD accession number 6G9T) as the template using the Modeller version 10.2.^19^ CAZ or AVI bound to CMY-185 was modeled by superimposition of the crystal structures of the AmpC^Ent385^ complex with CAZ (PBD accession number 6LC9) or AVI (PDB accession number 6LC8).^20^

### Data availability

Ec1, Ec2, and Ec3 whole genome sequencing data have been deposited in BioProject PRJNA924946. ST410 isolate WGS data from previous studies can be acquired in BioProject PRJNA836696 and BioProject PRJNA388450.^21, 22^

## Results and Discussion

### Clinical case

A male with a retroperitoneal liposarcoma was admitted for an abdominal tumor resection with reconstruction. One week post-operatively, he developed fever which was treated with piperacillin-tazobactam (TZP) for one week, at which time he was taken to surgery for a washout with drain placement. Two weeks after surgery, surgical drains grew a CAZ-resistant *E. coli* (baseline in **Table 1**), for which he was placed on ertapenem (ETP). After two weeks, the drain clogged and was repositioned, cultures obtained at that time grew carbapenem-resistant *E. coli* (Day 14 in **Table 1**). He was placed on CZA at this time with a planned 8-week treatment course. Near the end of therapy, the drain began bleeding, and he was transferred to the intensive care unit. A drain culture obtained at that time grew a CZA-resistant, carbapenem-susceptible *E. coli* (Day 65 in **Table 1**).

**Table 1:** Susceptibility Testing Results of Serial ST410 *E. coli* Isolates.

|  |  |  |  |
| --- | --- | --- | --- |
| **Antimicrobials** | **MICs (mg/L)** |  |  |
|  | ***E. coli* EC1**<br>**(Baseline)** | ***E. coli* EC2**<br>**(Day 14)** | ***E. coli* EC3**<br>**(Day 65)** |
| Aztreonam (ATM) | 16 (R) | >64 (R) | >64 (R) |
| Ceftazidime (CAZ) | >64 (R) | >64 (R) | >64 (R) |
| Ceftazidime-avibactam (CZA) | ND | 2 (S)<sup>b</sup> | >256 (R)<sup>b</sup> |
| Cefepime (FEP) | 2 (S) | >64 (R) | 32 (R) |
| Ertapenem (ETP) | <0.25 (S) | 16 (R) | 0.25 (S)<sup>a</sup> |
| Imipenem (IMI) | <0.25 (S) | 2 (I)<sup>a</sup> | 1.5 (I)<sup>a</sup> |
| Meropenem (MEM) | <0.25 (S) | 2 (I)<sup>a</sup> | 0.19 (S) |
| All testing was performed using Vitek2 unless otherwise indicated. ND, not done. |  |  |  |
| <sup>a</sup>Testing completed with Etest (bioMérieux). |  |  |  |

### Phenotypic and genotypic resistance of E. coli clinical isolates

The baseline *E. coli* isolate Ec1 was resistant to CAZ and ATM, whereas susceptible to cefepime (FEP) and carbapenems (**Table 1**). *E. coli* isolate Ec2, which was collected after two weeks of ETP therapy, was additionally resistant to FEP and ETP, and intermediate to imipenem (IMI) and meropenem (MEM), whereas it remained susceptible to CZA. *E. coli* isolate Ec3 was collected after a prolonged course of CZA and was highly resistant to this agent with an MIC of >256 mg/L. It was also resistant to other tested cephalosporins but susceptible to all carbapenems including ETP.

We determined that all three isolates belonged to ST410 based on *in silico* typing using WGS data. For the purpose of determining genetic relatedness of these serial isolates in the context of ST410 we have previously sequenced,^21, 22^ we created a maximum likelihood phylogenetic tree inferred from a recombination free, core genome alignment of 15 ST410 isolates collected from 10 patients (**Figure 2**). After accounting for recombination regions, there were 831 polymorphic sites with a median pairwise SNP distance of 210 (interquartile range = 52.5). Notably, Ec1, Ec2, and Ec3 had <10 pairwise SNP distances. The prominent genomic difference detected between Ec3 and its antecedent isolates (Ec1 and Ec2) is the acquisition of an IncIγ conjugative plasmid that is in association with the class C β-lactamase, pAmpC gene *bla*_CMY-185_ (Accession #: OQ297612.1) as highlighted in **Figure 2**. The predicted CMY-185 b-lactamase differed from the most commonly detected pAmpC in *E. coli, i.e*., the CMY-2 enzyme, by four amino acid substitutions (A114E, Q120K, V211S, N346Y). The other three ST410 IncIγ positive isolates in our population harbored *bla*_CMY-42_, which encodes for an enzyme that only differs by one amino acid (V211S) and showed moderate hydrolysis of CAZ compared to the CMY-2 enzyme.^23^ All four IncIγ positive isolates had 100% breadth of coverage to the pCMY42-035148 (RefSeq #: GCF_003268695.1) IncIγ plasmid. Interestingly, all *bla*_CMY_ positive isolates had a mutation in the *ftsI* gene which encodes PBP3, a key divisome target of cephalosporins and monobactams, where either a predicted ‘YRIN’ (*i.e*., YRIN N337N) or ‘K(P)YRI I336I’ duplication event occurred (**Figure 1**). Furthermore, Ec1, Ec2, and Ec3 had mutations in *ftsI* corresponding to three substitutions (Q227H + E349K + I532L). These FtsI substitutions have been associated with reduced susceptibility to CAZ and FEP,^24^ ATM-AVI in the presence of *bla*_CTX-M-15_,^25^ and associated with the acquisition of carbapenemases in ST410 backgrounds.^26^ A full table of all acquired AMR genes detected in Ec1 – Ec3 is presented in **Table 2**. These results suggest that *bla*_CMY-185_ may have developed through selective pressures that beget mutations in a *bla*_CMY-42_ gene harbored on an IncIγ conjugative plasmid.

**Figure 2.**
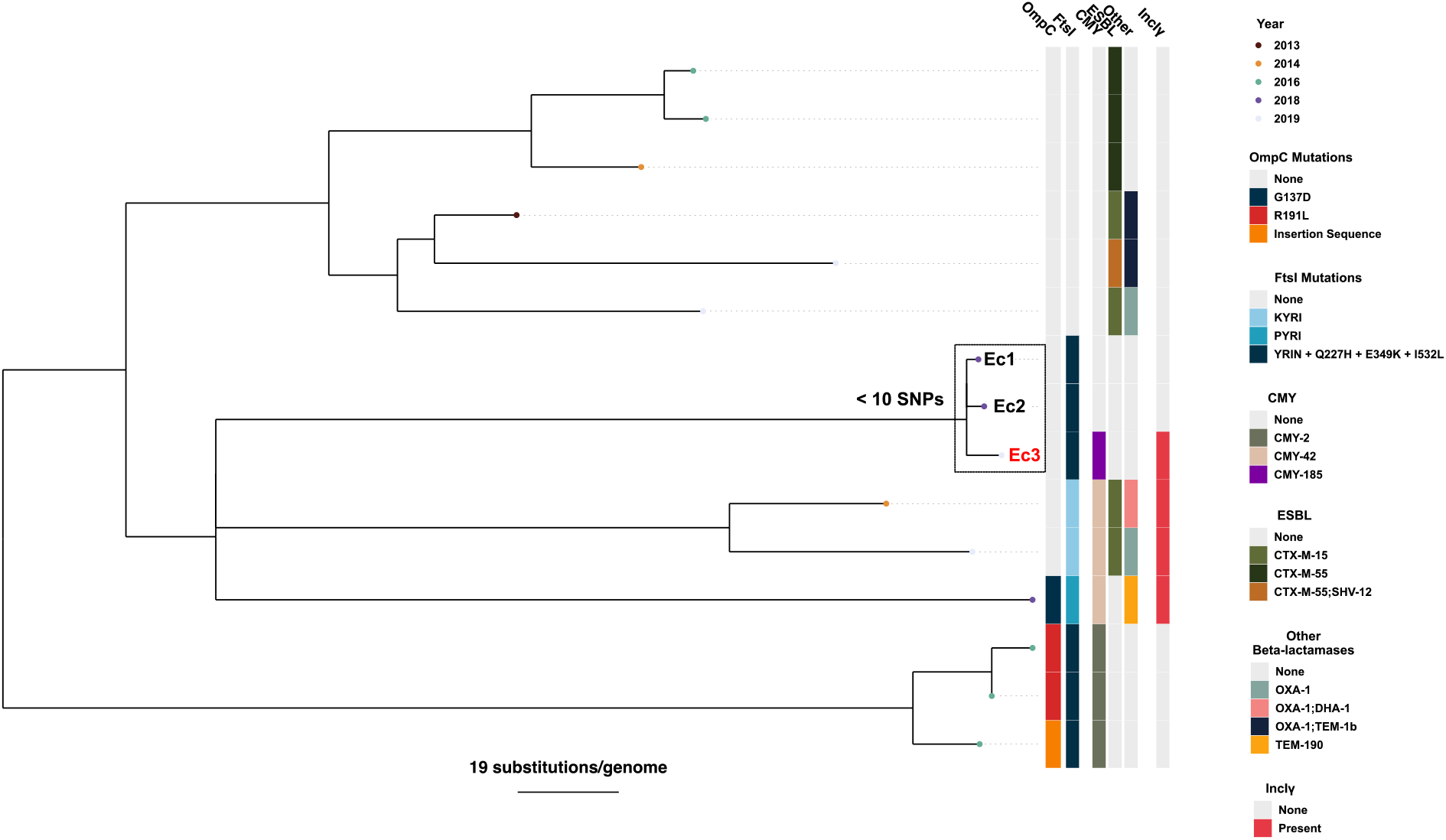
ST410 Population Structure with *bla*_CMY-185_-Harboring Strain. A total of 15 ST410 isolates from 10 patients have been identified through WGS from 2013 to 2019 locally. Ec1, Ec2, and Ec3 are indicated on the tree having <10 pairwise SNPs within this group. Ec3 is highlighted in red font with the novel *bla*_CMY-185_ variant (purple) indicated in the column. All IncIγ positive isolates had 100% breadth of coverage when aligning to pCMY42-035148 (RefSeq #: GCF_003268695.1). *ftsI* and *ompC* mutations that are predicted to contribute to reduced susceptibility to b-lactams based on previous studies ^24, 26^ are indicated in the legend.

**Table 2:** Acquired AMR Genes for Serial ST410 Isolates.

| **Table 2: Acquired AMR Genes for Serial ST410 Isolates** |  |  |  |
| --- | --- | --- | --- |
|  | *E. coli* Ec1 | *E. coli* Ec2 | *E. coli* Ec3 |
| **Acquired resistance** |  |  |  |
| β-lactams | NA | NA | *bla*<sub>CMY-185</sub> |
| Aminoglycosides | *aadA5, mphA,* | *aadA5, mphA,* | *aadA5, mphA,* |
| Folate synthase inhibitors | *dfrA17* | *dfrA17* | *dfrA17* |
| Fluoroquinolones | *qnrB7* | *qnrB7* | *qnrB7* |
| Sulfonamides | *sul1, sul2* | *sul1, sul2* | *sul1, sul2* |
| Tetracyclines | *tetB, tetD* | *tetB, tetD* | *tetB, tetD* |

### CZA resistance conferred by CMY-185

To determine the phenotype conferred by CMY-185, *bla*_CMY-185_ and *bla*_CMY-2_ were constitutively expressed in a cloning vector. CMY-185 conferred 256-fold higher MIC of CZA compared with CMY-2, confirming the role of the four substitutions in the CZA resistance observed in Ec3 (**Table 3**). Interestingly, it also conferred 16-fold higher MIC of cefiderocol (FDC), a siderophore cephalosporin recently approved for clinical use, in comparison with CMY-2. Co-resistance to CZA and FDC has been reported to occur in AmpC with genetic changes in the R2 region through increased hydrolytic efficiency of CAZ and FDC.^20, 27, 28^ However, none of the substitutions in CMY-185 are located in the R2 region, and both increased hydrolysis of CAZ and impaired inhibition by AVI appear to contribute to its CZA resistance phenotype (see discussion below).

**Table 3:** MICs of *E. coli* TOP10 strains carrying pBCSK(-) vectors that encode single, double, triple and quadruple (CMY-185) mutants of CMY-2.

|  |  |  |  |  |  |  |
| --- | --- | --- | --- | --- | --- | --- |
| **CMY-2 and its mutants** | **MICs (mg/L)** |  |  |  |  |  |
|  | **CAZ** | **CZA** | **FDC** | **FEP** | **ATM** | **MEM** |
| pBCSK(-) (control) | 0.25 | 0.12 | 0.06 | <0.03 | 0.12 | <0.03 |
| CMY-2 | 16 | 0.12 | 0.12 | 0.12 | 2 | <0.03 |
| Single mutants |  |  |  |  |  |  |
| A114E | 8 | 0.25 | 0.12 | 0.25 | 4 | <0.03 |
| Q120K | >32 | 0.5 | 0.25 | 0.25 | 4 | <0.03 |
| V211S | >32 | 0.25 | 0.12 | 0.12 | 4 | <0.03 |
| N346Y | 2 | 0.25 | 0.25 | <0.03 | 0.25 | <0.03 |
| Double mutants |  |  |  |  |  |  |
| A114E Q120K | 32 | 1 | 0.5 | 0.25 | 4 | <0.03 |
| A114E V211S | >32 | 0.25 | 0.25 | 0.12 | 16 | <0.03 |
| A114E N346Y | 16 | 8 | 0.5 | 2 | 2 | <0.03 |
| Q120K V211S | >32 | 0.25 | 0.5 | 0.12 | 8 | <0.03 |
| Q120K N346Y | 32 | 32 | 1 | 0.25 | 2 | <0.03 |
| V211S N346Y | 32 | 4 | 0.5 | 0.5 | 8 | <0.03 |
| Triple mutants |  |  |  |  |  |  |
| A114E Q120K V211S | >32 | 2 | 0.5 | 0.25 | 8 | <0.03 |
| A114E Q120K N346Y | 32 | 16 | 2 | 1 | 4 | <0.03 |
| A114E V211S N346Y | 32 | 16 | 1 | 0.5 | 16 | <0.03 |
| Q120K V211S N346Y | >32 | 16 | 1 | 0.25 | 8 | <0.03 |
| Quadruple mutant (CMY-185) |  |  |  |  |  |  |
| A114E Q120K V211S N346Y | >32 | 32 | 2 | 0.5 | 16 | <0.03 |
| CAZ, ceftazidime; CZA, ceftazidime-avibactam; FDC, cefiderocol; FEP, cefepime; ATM, aztreonam; MEM, meropenem. |  |  |  |  |  |  |

### Impact of amino acid substitutions observed in CMY-185

Given the presence of four amino acid substitutions in CMY-185, *bla*_CMY-2_ mutants with single, double, and triple amino acid substitutions contained in CMY-185 were constructed for every combination to decipher their roles in CZA resistance. None of the single amino acid mutant strains showed increased CZA MICs (**Table 3**). Rather, the N346Y mutant strain appeared to be impaired in its hydrolytic activity of CAZ and ATM, with 8-fold decrease in MICs compared with the strain producing CMY-2. In contrast, all three double mutant strains containing the N346Y substitution showed CZA MICs that were 32 to 256-fold higher than that of the CMY-2-producing strain, without major differences in CAZ MICs. These observations suggested that secondary substitutions were required in addition to N346Y for the enzymes to confer CZA resistance, most likely due to impaired inhibition by AVI. This trend was extended to triple mutations, where the three triple mutant strains containing N346Y showed a CZA MIC of 16 mg/L, 128-fold increase compared with CMY-2. The triple mutant strain from which N346Y was absent (A114E, Q120K, V211S) had a CZA MIC of 2 mg/L, which was 16-fold higher than that of CMY-2. On balance, the quadruple mutant (CMY-185) had the highest MICs across the tested cephalosporins (CAZ, CZA, FDC) except for FEP. All the mutants remained susceptible to MEM.

### Molecular modelling

To visualize the substitution sites, we generated a homology-model structure of CMY-185 (**Figure 3A**). The model structure of CMY-185 showed that the A114E substitution is located at the inner-molecule assembled by the hydrophobic contact with W100, I104, L109, L117 and W138. The Q120K, V211S and N346Y substitutions are located at the molecular surface where the substrate is bound. The N346 residue is a significant residue for interacting with the cephalosporins or AVI and make hydrogen bonds with carboxyl group at 4-position of the cephalosporin ring and the amide group at 2-position of AVI (**Figure 3DE**). Compain et al. reported that the N346Y substitutions in class C b-lactamases AmpC_cloacae_, PDC-5 and DHA-1 bring about large decreases in carbamoylation efficacy of AVI due to the steric hindrance by the bulky side chain of tyrosine (**Figure 3BC**).^6^ In contrast, the impact of the N346Y substitution on the MIC of CAZ depends on the enzyme. The N346Y mutants of AmpC_cloacae_ or PDC-5 showed moderate hydrolysis of CAZ, while the N346Y mutant of DHA-1 showed 4-fold decrease in the MIC of CAZ.^6^ The single N346Y mutant of CMY-2 showed 8-fold decrease in the MIC of CAZ compared with a wild type CMY-2 in this study. These results suggest that the reduction of the inhibition of AVI appears to be a major cause for the reduced susceptibility to CZA by the N346Y substitution in CMY-2. However, the secondary substitutions were required in addition to the N346Y substitution to confer CZA resistance in CMY-2. Given the concerning susceptibility patterns, the additional A114E, Q120K and V211S substitutions appear to restore the resistance to CAZ diminished by the N346Y substitution and further to FDC, which contains a bulkier R2 side chain. Further studies are needed to understand the molecular mechanism for CZA or FDC resistance conferred by the combination of N346Y substitution with A114E, Q120K and V211S substitutions.

**Figure 3.**
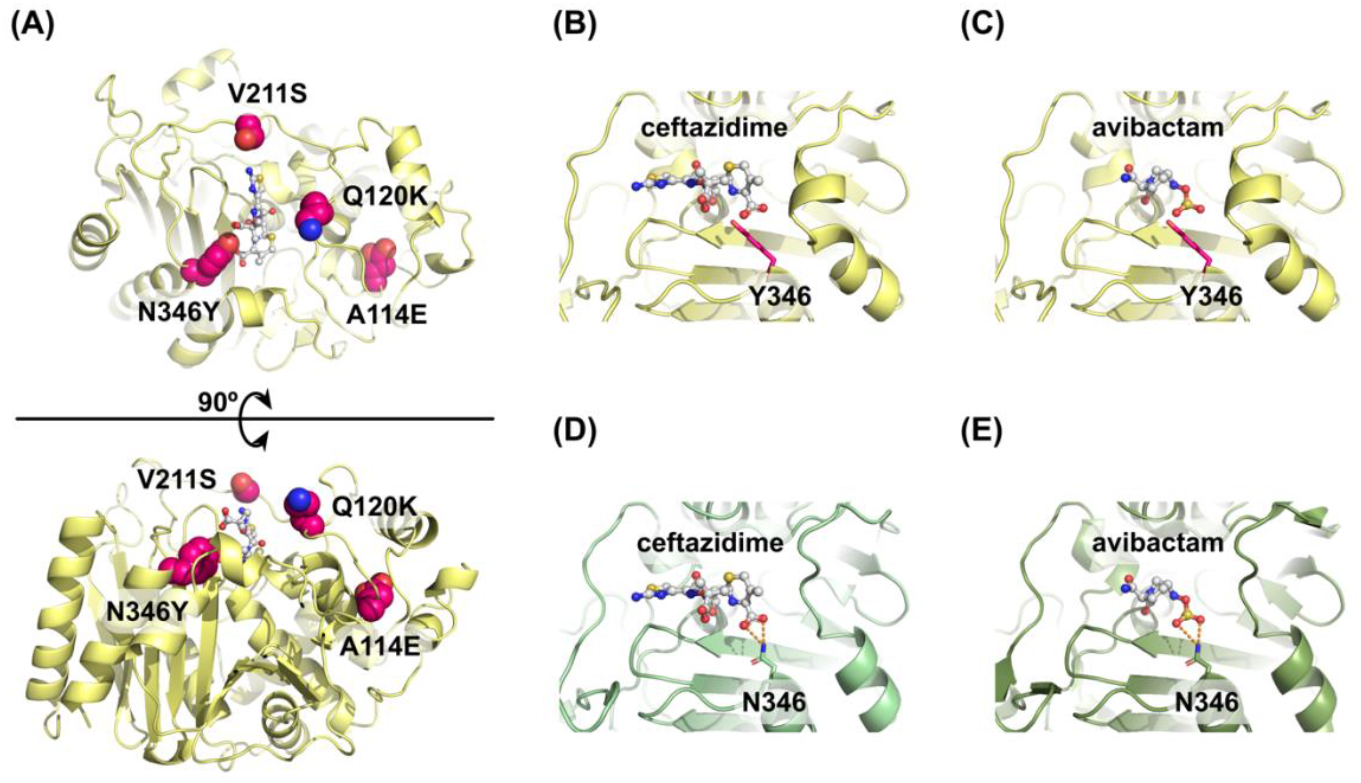
Model structure of CMY-185. (A) Overall structure of CMY-185. The substitution residues are shown as CPK representations colored in magenta. To clarify the substrate binding site, the ceftazidime molecule is shown as a white ball-and-stick representation. (B, C) Close-up views of the model structures of the CMY-185 complex with ceftazidime (B) or avibactam (C). (D, E) Close-up views of the crystal structures of the AmpC^Ent385^ complex with ceftazidime (D, PDB accession number 6LC9) or avibactam (E, PDB accession number 6LC8). Hydrogen bonds are represented as orange dashed lines.

### Conclusion

Our case illustrates how CMY-2, the most common pAmpC b-lactamase, may mutate upon specific selective pressure and result in CZA resistance during clinical treatment. Additionally, this study also highlights a unique enzymatic feature where secondary substitutions complement the b-lactam degradations diminished by trading off the substitution to escape from the inhibitor. Given the widespread and transferrable nature of CMY and other pAmpC enzymes in *E. coli* and other *Enterobacterales* species, this finding raises concern for additional cases of resistance with increasing usage of CZA.

## Transparency Declarations

The authors declare no conflicts of interest.

## Author contributions

W.C.S.: data acquisition, WGS, computational analyses, manuscript drafting; B.T.E.: data abstraction, EHR review, manuscript preparation; J.B.: data abstraction, EHR review, manuscript preparation; S.L.A.: data abstraction, data curation, methodology, manuscript preparation; W.C.B.: mutant construction; C.L.M.: microbiology; A.K.: genomic analysis; M.M.B.: data acquirement, antimicrobial susceptibility testing; P.S.: antimicrobial susceptibility testing, isolate characterization; A.K.: molecular modeling, manuscript preparation; R.K.S.: data curation, manuscript drafting; S.A.S.: funding acquisition, supervision, data acquirement, manuscript drafting; Y.D.: funding acquisition, supervision, conceptualization, data analysis, manuscript drafting/editing

